# Shared understanding is correlated with shared neural responses in the default mode network

**DOI:** 10.1101/231019

**Authors:** Mai Nguyen, Tamara Vanderwal, Uri Hasson

## Abstract

Humans have a striking ability to infer meaning from even the sparsest and most abstract forms of narratives. At the same time, flexibility in the form of a narrative is matched by inherent ambiguity in the interpretation of it. How does the brain represent subtle, idiosyncratic differences in the interpretation of abstract and ambiguous narratives? In this fMRI study, we scanned subjects watching a 7-min original animation that depicts a complex narrative through the movement of simple geometric shapes. We additionally scanned two separate groups listening to concrete verbal descriptions of either the social narrative or the physical properties of the movie. After scanning, all subjects freely recalled their interpretation of the stimulus. Using an intersubject representational similarity analysis, we compared the similarity of narrative interpretation across subjects, as measured using text analysis, with the similarity of neural responses, as measured using intersubject correlation (ISC). We found that the more similar two people’s interpretations of the abstract shape movie were, the more similar their neural responses were in the default mode network (DMN). Moreover, these shared responses were modality invariant: despite vast differences in stimulus properties, we found that the shapes movie and the verbal interpretation of the movie elicited shared responses in linguistic areas and a subset of the DMN when subjects shared interpretations. Together, these results suggest that the DMN is not only sensitive to subtle individual differences in narrative interpretation, but also resilient to large differences in the modality of the narrative.

**Significance statement:** The same narrative can be both communicated in different ways and interpreted in different ways. How are subtle, idiosyncratic differences in the interpretation of complex narratives presented in different forms represented in the brain? In this fMRI study, we show that the more similarly two people interpreted an ambiguous animation composed of moving shapes, the more similar their neural responses were in the Default Mode Network. In addition, by presenting the same narrative in a different form, we found shared responses across modalities when subjects shared interpretations despite the vast differences in modality of the stimuli. Together, these results suggest that the DMN is at once sensitive to individual differences in narrative interpretation and resilient to differences narrative modality.

## Introduction

Human communication is remarkably flexible, and the same narrative can be communicated in many different ways, from words to pictures and even via the motion of simple geometric shapes (1). At the same time, narratives can be inherently ambiguous, leading to differences in interpretation. How are these kinds of subtle and idiosyncratic differences in the interpretation of complex, ambiguous events—which may be communicated in different forms—represented in the brain?

Previous work using intersubject analyses have shown that complex, temporally-extended narratives elicit shared neural responses across subjects in a network of high-order brain regions, including temporal parietal junction (TPJ), angular gyrus, temporal poles, posterior medial cortex (PMC), and medial prefrontal cortex (mPFC) (2–8). The correlated responses in these high-level regions, which largely overlap with the Default Mode Network (DMN), seem to be driven mainly by the content of the narrative, independent of the form or modality in which it is communicated. For example, researchers have identified shared neural responses between subjects who listen versus read the same narrative (9), between subjects who watch an audiovisual movie versus listen to a verbal description of the movie (10–12), and between subjects who listen to a Russian versus English translation of the same narrative (13).

Shared neural responses in these studies are thought to reflect shared perception and subsequent interpretation of the stimuli. A recent study supports this view: researchers in this study manipulated the interpretation of a narrative by providing two separate groups with different contextualizing information, resulting in two coherent, but opposing, interpretations of the same narrative. The researchers found group-level differences in the similarity of neural responses in the DMN: similarity was greater within the subjects who shared the same interpretation of the narrative than between subjects who had opposing interpretations (14).

In these previous studies, however, interpretation of the narrative was either uniform across subjects or directly manipulated. As a result, all subjects within a group share the same interpretation of the narrative. In real-life, narratives can be ambiguous, leading to many different interpretations across subjects. To what extent then, will the spontaneous and unguided interpretation of an ambiguous, abstract narrative covary with the level of shared neural responses across subjects? Based on previous work, we first predict that the more similar the interpretation of an abstract movie across individuals, the more similar their neural responses in the DMN will be. In addition, we predict that because the DMN has been shown to represent narrative content independent of modality, we should observe a similar relationship between similarity of interpretation and similarity of neural response even when narrative is presented in two different modalities: moving geometric shapes versus spoken words.

To test the first prediction, we created an abstract, ambiguous seven-minute long animated sequence in which two triangles, a circle, a square, and a group of rectangles interact, following the classic work of Heider & Simmel (1). Work using animated shape movies have shown that the majority of subjects attribute animacy to the movement of these simple geometric shapes (1, 15–17). However, subjects also typically show substantial variance in their interpretation of the actual narrative (1, 15, 18, 19). Unique to this study, the animation is significantly longer than the typical shapes movie (e.g. 7 minutes versus 10-30 seconds), contains numerous characters with different relationships (e.g. parent and child, friends, antagonists), and a high-level narrative arc with complex and more abstract social and psychological events (see SI M1 for animation).

We scanned subjects in fMRI while they watched the animation and then asked them to describe their detailed interpretation of the narrative (Fig 1A,B). Using text analysis tools, we assessed how similar each subject’s interpretation of the animation was to every other subject’s interpretation. Using intersubject analyses, we also measured the similarity of each subject’s neural response to every other subject’s in each brain area. Finally, we correlated the interpretation similarity matrix with the neural similarity matrix using a Representational Similarity Analysis (RSA) (20) to test whether shared interpretation predict shared neural response (Fig 2).

**Fig. 1.**
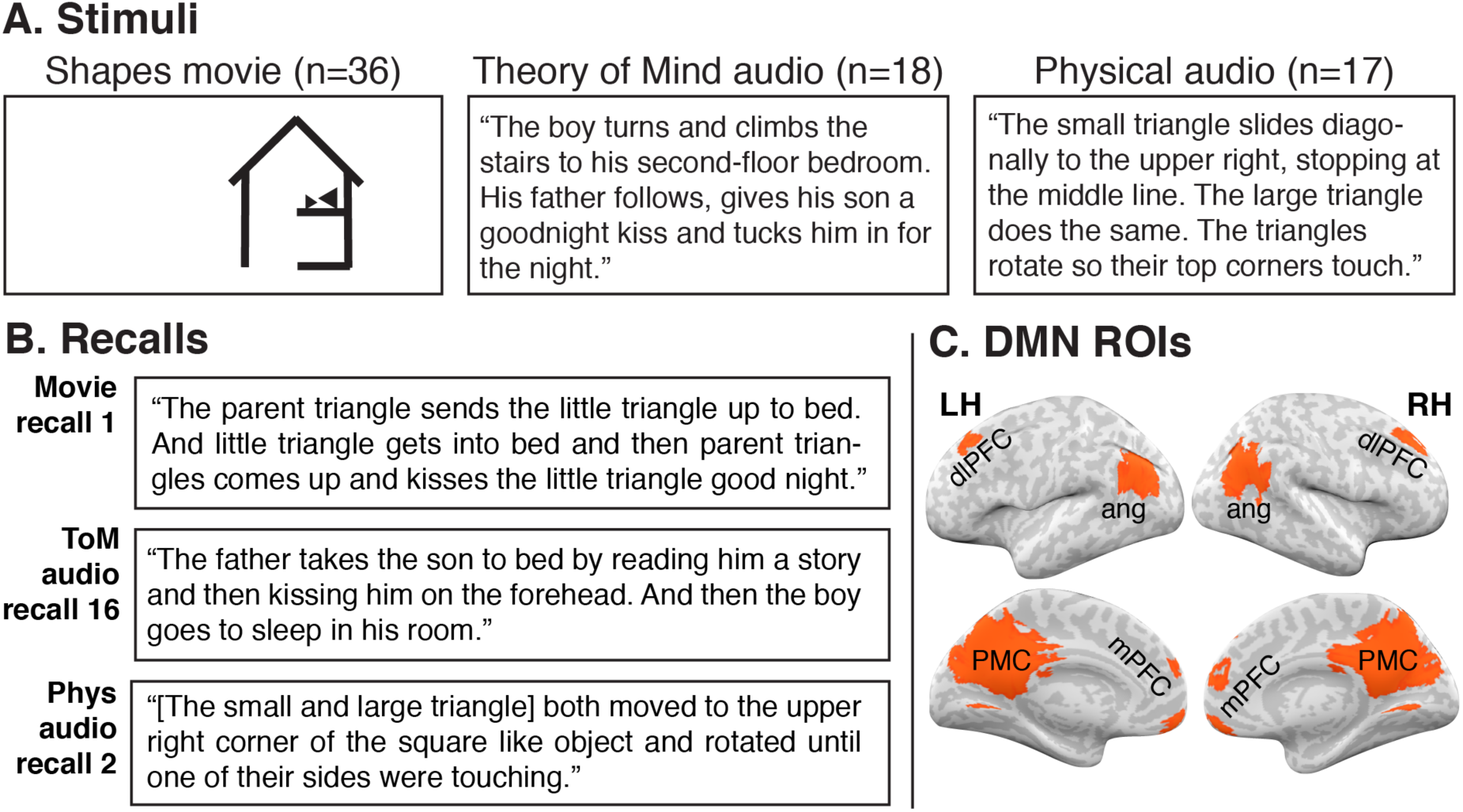
Design. (A) While being scanned in fMRI, subjects watched a short animation made of moving shapes (Movie), listened to an audio version of the movie narrative (Theory of Mind Audio), or listened to an audio description of the moving shapes in the animation (Physical Audio). (B) Following stimulus presentation, subjects retold the story they heard. Excerpts from a subject in each group are shown here. Recalls were assessed for similarity within and across stimulus groups using latent semantic analysis (LSA). (C) Default mode network ROIs defined using independent data. PMC = posterior medial cortex, mPFC = medial prefrontal cortex, ang = angular gyrus, dlPFC = dorsolateral prefrontal cortex.

**Fig. 2.**
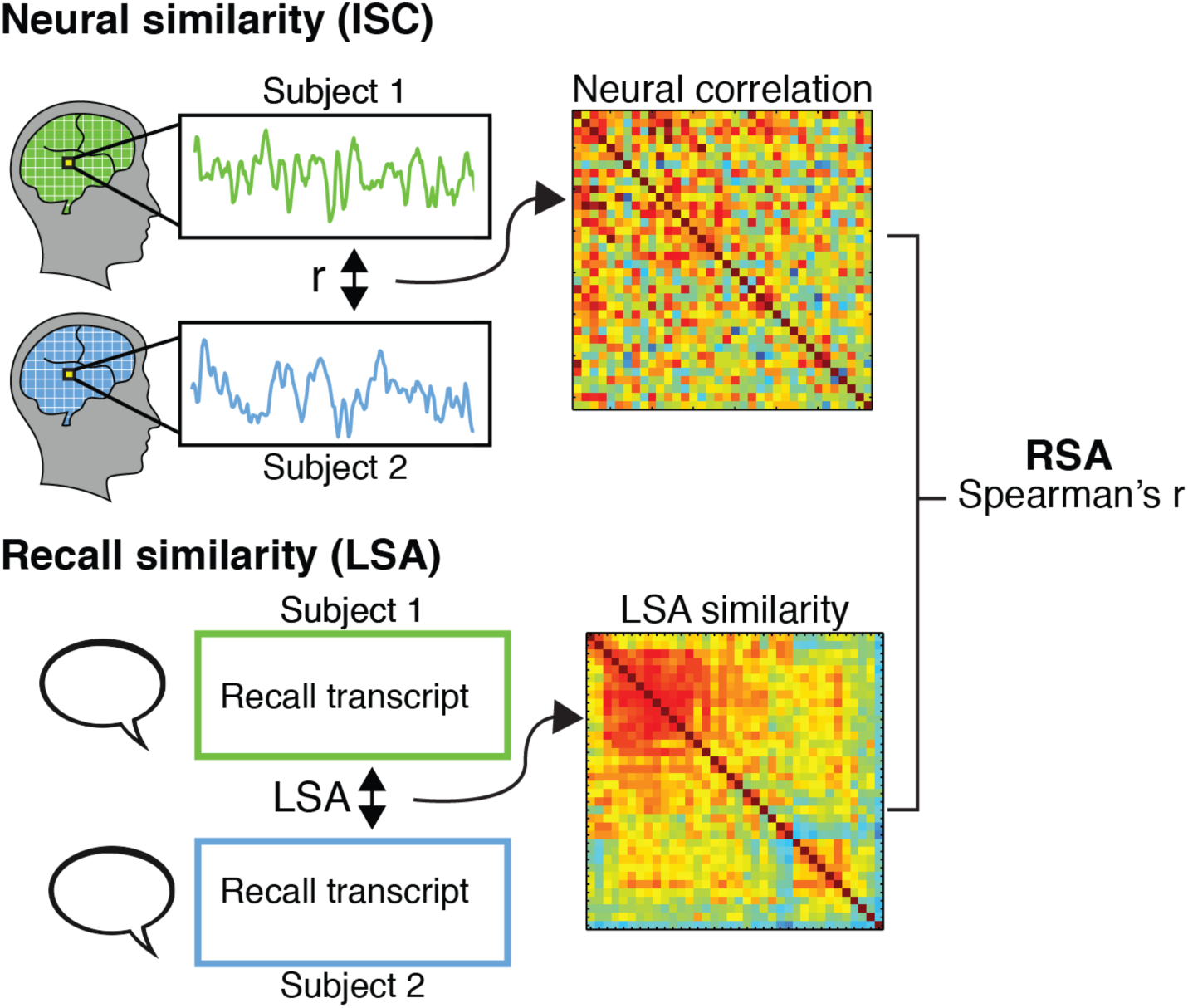
RSA. An intersubject representational similarity analysis (RSA) was conducted to identify regions of the brain where greater similarity in narrative interpretation was correlated with greater neural similarity. To measure neural similarity, each subject’s response timecourse was correlated with every other subject’s timecourse in every voxels. To measure interpretation similarity, we used Latent Semantic Analysis (LSA) to measure semantic similarity between the recalls of every pair of subjects. We then calculated the spearman r between the neural and recall similarity matrices in all voxels of the brain. Statistical significance was assessed using a permutation test and then FDR corrected for multiple comparisons. ISC = intersubject correlation, LSA = latent semantic analysis, RSA = representational similarity analysis.

To test the second prediction that the same narrative presented in different forms should elicit shared neural responses as long as the interpretation is the same, we additionally scanned two separate groups of subjects listening to two explicit verbal descriptions of the animation. In one condition, subjects listened to a possible social interpretation of the animation given by the animators (e.g. “the father tucked his son in for a night”). In the second condition, subjects listened to a verbal description of the physical motion of the shapes in the animation (e.g. “the large triangle rotated until it overlapped with the small triangle”) (Fig 1A). We then tested for shared interpretation across stimulus modality by both correlating interpretation similarity with neural similarity across modalities using RSA (Fig 2) and by directly correlating neural responses between the movie group and the two audio groups using ISC.

We found that the more similar two people’s spontaneous and unguided interpretation of the animation were, the more similar were their neural response in the DMN. Moreover, we found modality invariant shared neural responses in linguistic areas, bilateral angular gyrus, and PMC between subjects who watched the animation and subjects who listened to the linguistic based interpretation of the animation, but only when the movie subjects shared the interpretation of the animator. Finally, we found that, irrespective of interpretation, there were shared neural responses between the movie and the description of the physical motion in visual areas and superior parietal lobule. Together, these results suggest that the default mode network is at once incredibly sensitive to subtle, individual differences in narrative interpretation and remarkably resilient to vast differences in the form of narrative communication.

## Results

We compared the behavioral and neural responses within and between three different groups to identify areas of the brain that represent shared understanding of ambiguous narratives over time. The “Movie” group was scanned in fMRI while they watched a 7-min ambiguous animation that told a complex social narrative using only the movement of simple geometric shapes. The “Theory of Mind (ToM) Audio” group was scanned while listening to a verbal description of the social interactions in the animation as interpreted by the animator (e.g. the father tucks his son into bed) while the “Physical (Phys) Audio” group listened to a description of the physical movements of the shapes in the animation (e.g. the corner of a large triangle touches a smaller triangle) (Fig. 1). The audio conditions were time-locked to the animation, so that events across all three stimuli were temporally correlated.

### Behavioral results

#### Variance in shared interpretation across subjects

We used Latent Semantic Analysis (LSA) (21), to measure the similarity of recall between each pair of subjects within the three groups and between the movie group and the two audio groups (see methods for more detail). We then used agglomerative hierarchical clustering with complete linkage to order subjects based on the similarity of their interpretation to each other (Fig. 3). The mean LSA similarity within the Movie Group was 0.619 (std = 0.125, range=0.26-0. 89; Fig. 3,top left square), the ToM Audio group 0.852 (std = 0.045, range=0.70-0.93; Fig. 3, middle square), and the Phys Audio group 0.638 (std = 0.94, range=0.36-0.84; Fig 3, bottom right square). The recalls in the Movie group were significantly less similar to one another than the recalls in the ToM Audio group (t(781) = 22.7, p<.001). There was no difference in similarity within the Movie group and the Phys Audio group (t(764) = 1.63, p>.05).

**Fig. 3.**
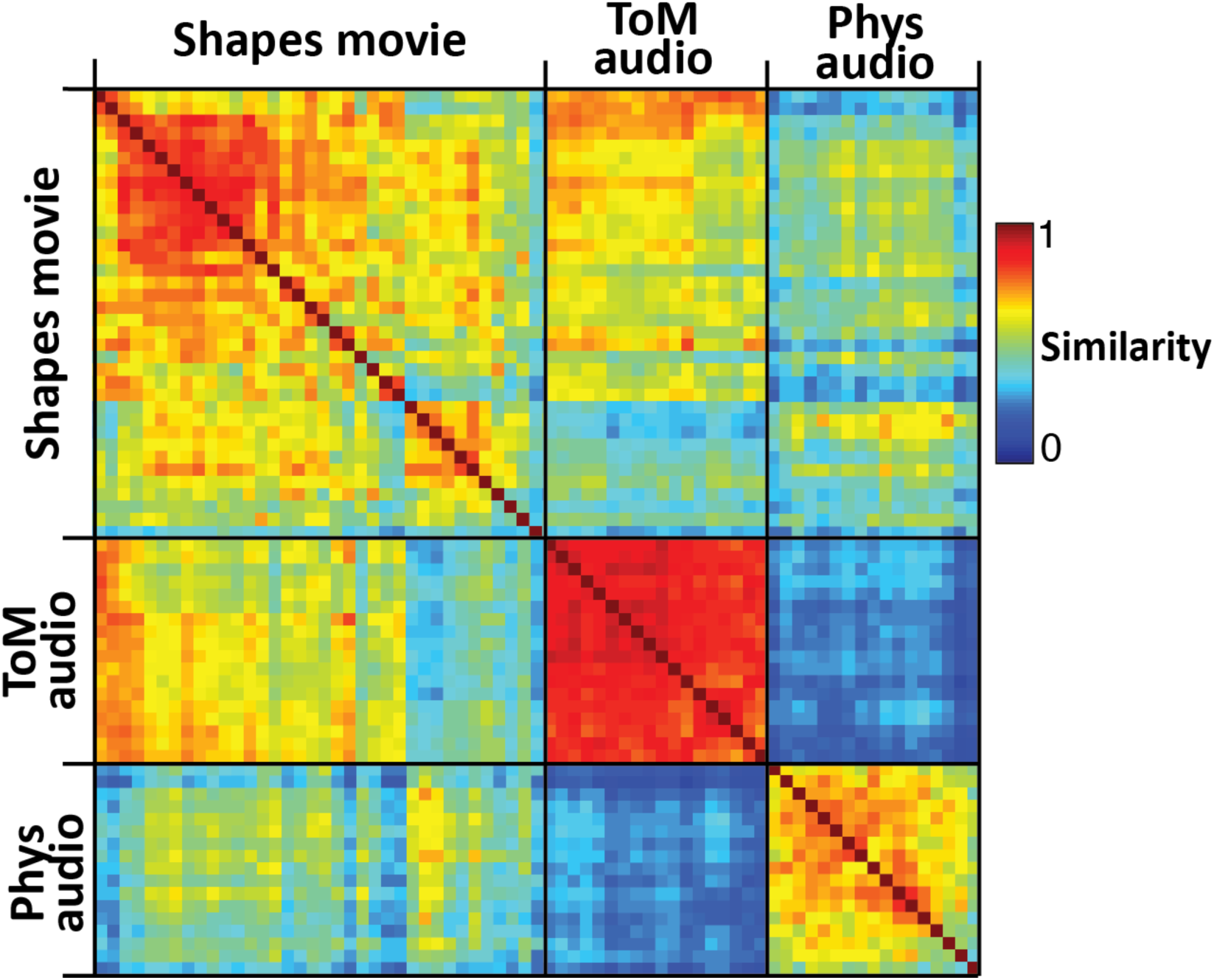
Behavioral results. Recall similarity between each pair of subjects in each group was assessed using Latent Semantic Analysis. Significantly greater intersubject similarity was observed in ToM Audio group than Movie group (t=22.7, p<.001); there was no difference in recall similarity within the Movie group and Phys Audio group (p>.05). Between groups, there was more recall similarity between Movie and ToM audio subjects than between Movie and Phys audio subjects (t=17.0, p<.001).

Between groups, the average LSA similarity between the Movie group and the ToM Audio group was 0.545 (std=0.125, range=0.26-0.8). Between the Movie group and the Phys Audio group, the average LSA similarity was 0.434 (std=0.105, range=0.18-0.72). The Movie recalls were significantly more similar to those in the ToM Audio group than those in the Phys Audio group (t(1258)=17.0, p<.001).

### Neural results

#### RSA: Interpretation similarity is correlated with neural similarity

We hypothesized that greater similarity in the interpretation of an ambiguous animation of geometric shapes will be reflected in greater neural similarity across subjects in high-order areas which encode the animation’s narrative. To test this prediction, we conducted an intersubject Representational Similarity Analysis (RSA) (20), over the entire brain: we correlated the timecourse of neural activation between pairs of subjects within or between groups for every voxel in the brain, and then correlated these neural correlations with the LSA similarity between each pair of subjects (Fig. 2, see methods for details).

#### RSA in Movie group

We first compared the neural and recall similarity in the Movie group in a whole-brain, voxel-wise RSA. We found that the level of recall similarity was correlated with the level of neural similarity in PMC, right angular gyrus, right anterior STS, left supramarginal gyrus, bilateral DMPFC, and bilateral DLPFC (q<.05, FDR corrected; Fig. 4A). The same correlations were also measured in six independently defined ROIs of the Default Mode Network (Fig. 4B).

**Fig. 4.**
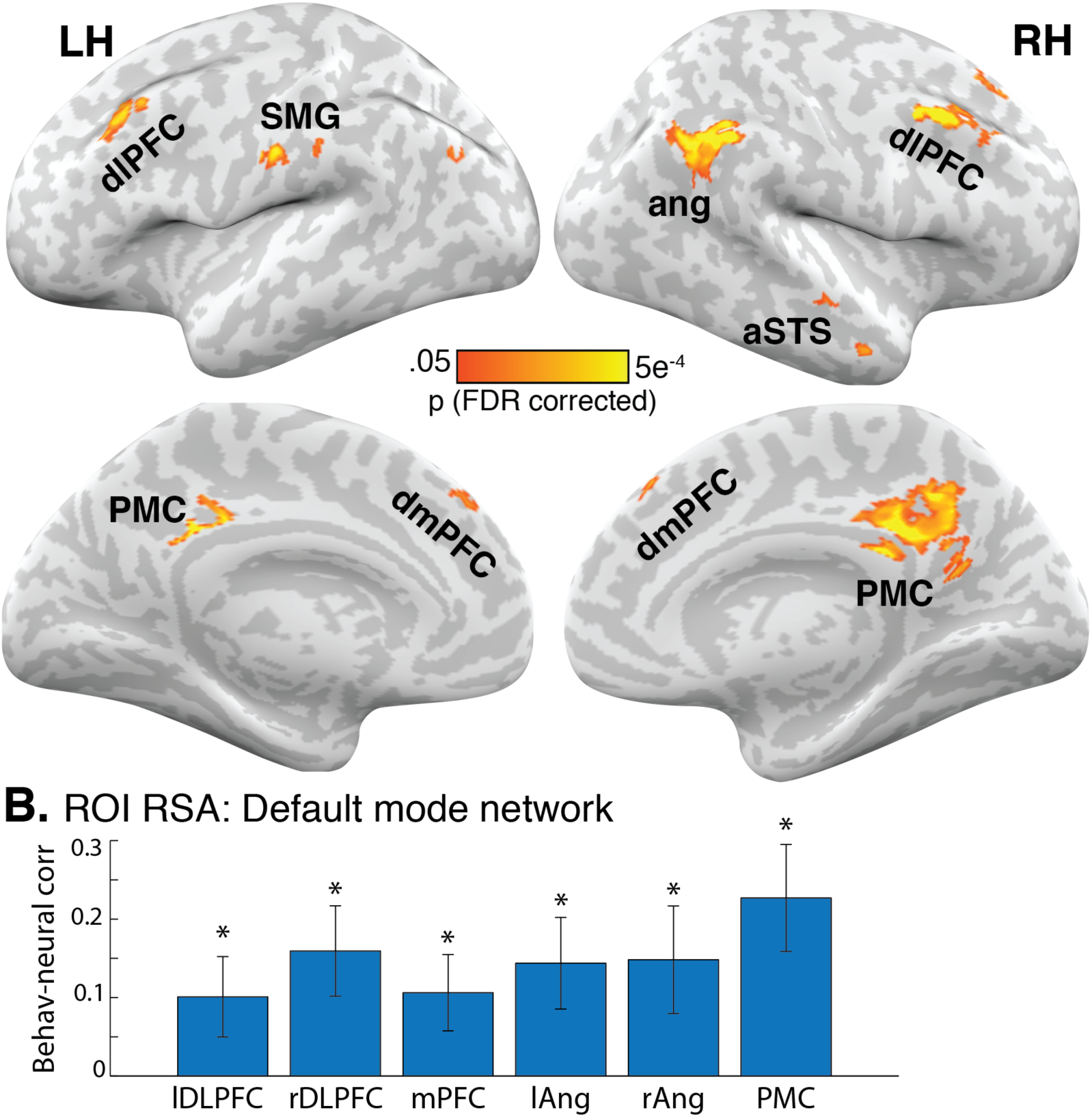
Shared interpretation, shared neural response. (A) Neural similarity in each voxel, measured by correlating response timecourses between every pair of subjects in the Movie group, was correlated with recall similarity, measured using Latent Semantic Analysis on the recalls of every pair of subjects in the Movie group. Neural similarity and recall similarity were significantly correlated with each other, as assessed using a permutation test, in PMC, dmPFC, dlPFC, left SMG, right angular gyrus, and right anterior STS. (B) RSA was also conducted on the average response of ROIs in the default mode network. All ROIs were significant in the movie group analyses (q<.05, FDR corrected). dlPFC = dorsolateral prefrontal cortex, SMG = supramarginal gyrus, ang = angular gyrus, aSTS = anterior superior temporal sulcus, PMC = posterior medial cortex, dmPFC = dorsomedial prefrontal cortex. * q< .05, FDR corrected. Error bars are SEM.

#### *RSA in Audio* groups

By design, the interpretations of the two audio recording were explicit and unambiguous. Due to the lack of variability in interpretation across subjects (Fig. 3), no significant voxels were identified by comparing the variance in neural and recall similarity in the ToM Audio group or the Phys Audio group in whole-brain analyses.

#### RSA across Movie and Audio groups

We additionally compared interpretation similarity and neural similarity between the Movie group and the two Audio groups. In the whole-brain analysis, no voxels passed significance testing in the Movie-ToM Audio comparison, although using a lower threshold revealed largely the same voxels as in the Movie, within-group RSA. The ROI analysis did reveal significant correlations between neural and recall similarity between the two groups in right DLPFC and PMC (q<.05, FDR corrected; Fig. 5). Finally, no voxels or ROIs passed thresholding for the Movie and Phys Audio comparison (q<.05, FDR corrected; Fig. 5).

**Fig. 5.**
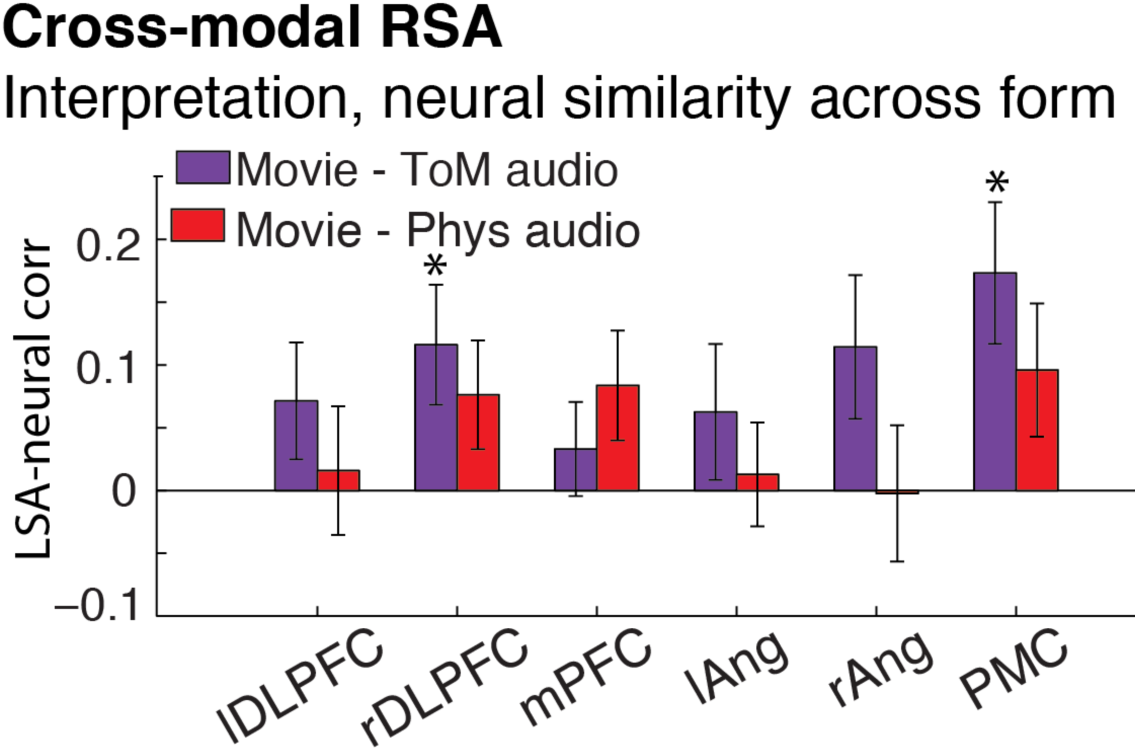
Greater neural similarity was correlated with greater interpretation similarity between the Movie group and the ToM Audio group in right DLPFC and PMC (purple bars). There was no relationship between neural and interpretation similarity between the Movie group and the Phys Aud group (red bars). dlPFC = dorsolateral prefrontal cortex, SMG = supramarginal gyrus, ang = angular gyrus, aSTS = anterior superior temporal sulcus, PMC = posterior medial cortex, dmPFC = dorsomedial prefrontal cortex. * q< .05, FDR corrected. Error bars are SEM.

#### ISC: shared response across animation and audios

In addition to conducting RSA, which is a second-order analysis based on the similarity of subjects to one another, we used intersubject correlation (ISC) (22) to directly compare the timecourse of neural responses between Movie subjects and either the ToM Audio subjects or the Phys Audio subjects. For each comparison (Movie-ToM Audio or Movie-Phys Audio), the movie subjects were split into “similar-to-audio” and “dissimilar-to-audio” groups based on their mean LSA recall similarity to the respective audio subjects. ISC was then calculated by correlating each Movie subject’s timecourse of activity within the group with the average timecourse of ToM Audio or Phys Audio subjects in every voxel (see methods for details).

#### Movie-ToM Audio ISC

The ToM Audio subjects and the 18 “similar-to-ToM-Audio” Movie subjects showed correlated neural responses in linguistic areas, including pSTS and ITG, and high-level areas including bilateral angular gyrus and PMC (p<.05, FWER corrected; Fig 6A, left). In contrast, the ToM Audio subjects and the 18 “dissimilar-to-ToM-Audio” Movie subjects only showed significant neural similarity in a small cluster of voxels in right angular gyrus (p<.05, FWER corrected; Fig 6A, right). In ROI analyses of the DMN, ISC between similar Movie subjects and ToM Audio subjects was significantly greater than between dissimilar Movie subjects and ToM Audio subjects in right DLPFC and PMC (q<.05, FDR corrected, Fig 6B).

**Fig. 6.**
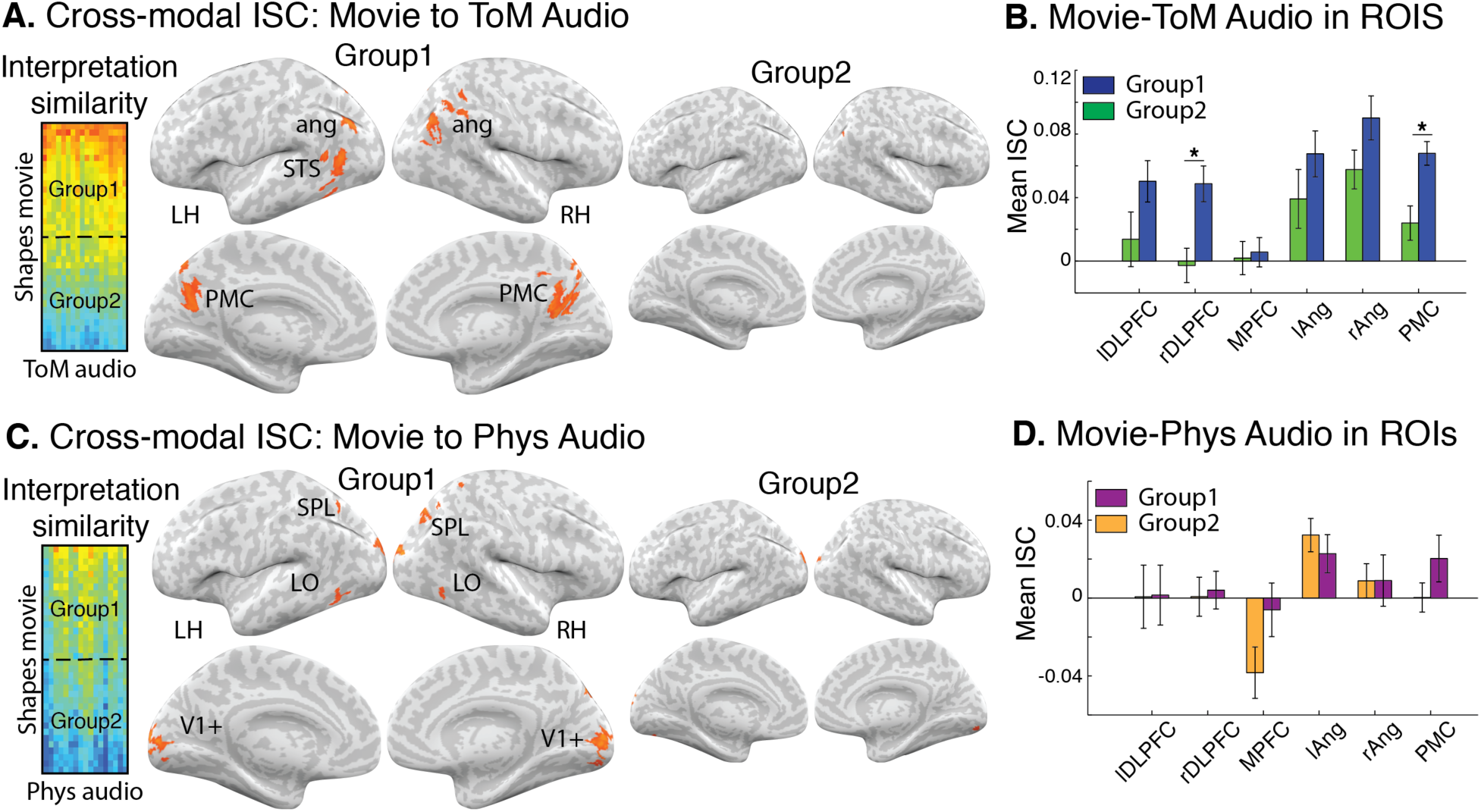
ISC between Movie and Audio groups. (A). Movie subjects who shared narrative interpretation with the ToM Audio subjects also showed correlated neural responses with the ToM Audio subjects in linguistic regions, bilateral angular gyrus and PMC (left brains). Movie subjects who did not share narrative interpretations with the ToM Audio group did not show correlated neural responses with ToM Audio subjects (right brains). (C). Movie subjects who shared descriptive content with the Phys Audio subjects showed correlated neural responses with the Phys Audio subjects in early visual cortex, LO, and SPL. Movie subjects who did not share descriptions only showed similar responses in early visual cortex. (B, D). ROI analysis using ISC show similar pattern of results. dlPFC = dorsolateral prefrontal cortex, SMG = supramarginal gyrus, ang = angular gyrus, aSTS = anterior superior temporal sulcus, PMC = posterior medial cortex, dmPFC = dorsomedial prefrontal cortex. * q< .05, FDR corrected. Error bars are SEM.

#### Movie-Phys Audio ISC

The Phys Audio subjects and the 18 “similar-to-Phys-Audio” Movie subjects showed correlated neural responses in bilateral early visual cortex, visual area LO, and superior parietal lobule (Fig 6C, left). The 18 “dissimilar-to-Phys-Audio” Movie subjects and Phys Audio subjects only showed similar neural responses in a small part of early visual cortex (Fig 6C, right). There were no significant differences between groups in ROI analyses (Fig 6D). Together, these results suggest that the DMN tracked the ToM interpretation and not the non-social interpretation of the narrative.

## Discussion

The same narrative or event can be interpreted in many different ways: for example, football rivals watching the same game both perceive that the opposing team commits more fouls even though they are watching the same game (23). However, the same narrative can also be communicated effectively using vastly different forms, including written or spoken word, sign language, and even moving abstract shapes, as in the present work.

We found that the more similarly two people interpreted the social events depicted in an ambiguous animation composed of moving shapes, the more similar their neural responses were in the DMN. In addition, we found that subjects who shared interpretation of the moving shapes not only had similar neural responses to each other, but also to subjects who heard a verbal description of the same interpretation of the narrative. The modality-invariant representation of shared interpretation occurs even despite the vast differences between moving geometric shapes and spoken words. In contrast, verbal descriptions of the physical aspects of animation elicited shared neural responses with the movie viewing in only visual cortex and superior parietal lobule.

This work is the first to show that the DMN, which includes medial cortex, angular gyrus, medial prefrontal cortex (24), can discriminate between idiosyncratic, spontaneous differences in the narrative interpretation of moving geometric shapes. These results are consistent with previous studies that directly manipulated interpretation by directing attention to different aspects of a spoken narrative (25), changing perspective (26), or biasing with contextual information (14). Unlike these previous results, however, in the present work, we do not manipulate the interpretation of the animation into discrete experimental groups. Rather, we let subjects freely attribute intentions to the motion of simple geometric shapes, which spontaneously led to the creation of complex social narratives in subjects' minds as expressed in their post-viewing descriptions of the animation. We then show that the subtle individual differences in rich narrative interpretation are reflected in individual differences in the neural responses of high-level regions.

Notably, these regions have also previously been implicated in theory of mind research, including the attribution of animacy to geometric shapes (27). In the typical study, researchers presented simple shape movies that either had animate motion (e.g. chasing, kicking) or non-animate motion (e.g. random motion). Greater activation was found during animate than inanimate movies in regions of DMN (28–30). By using a novel shapes animation that is unique for its length (7 mins vs the typical 10-30 seconds) and complex narrative arc, as well as the large number of interacting characters with different relationships (parent and child, friends, antagonists), the present work builds on previous findings by showing that these areas not only respond preferentially to animate films, but can discriminate between subtle differences in interpretations of the social interactions.

This work is also the first to directly show shared neural representations of a moving shapes animation and a verbal description of the same narrative in left pSTS, left ITG, PMC, and bilateral angular gyrus, speaking to the modality invariance of the DMN. Previous work has shown that parts of the DMN respond similarly to individual presented words versus images of the same item (e.g. 30–33); spoken versus written sentences, paragraphs or narratives (8, 36–38), and audio-visual versus spoken narratives (10–12). However, the present work is the first to show that despite vast differences in stimulus properties, sparse and abstract stimuli (like triangles and squares) elicit similar neural responses to explicit verbal narratives as long as both stimuli induce similar interpretations of the stimuli.

When narrative content was shared across the animation and verbal interpretation of the animation in areas of the DMN, we found that the physical, visual content of the animation was shared with a verbal description of the shapes in visual cortex (V1+, VO, LO) and bilateral superior parietal lobule (Fig. 4B). Shared neural activity in both regions is consistent with mental imagery during the Physical Audio task. While listening, subjects may have been mentally visualizing moving shapes and the spatial relations among them. Some previous work has demonstrated early visual cortex activation during mental imagery (39–42), while SPL has been implicated in mental rotation of shapes (43, 44). In addition, SPL and the neighboring intraparietal sulcus are involved in processing geometric and mathematical information such as numerosity (45, 46), geometric shape terms (47), and mathematical visual-spatial representations more broadly (48). Shared, cross-modal activity in SPL may thus arise from shared geometric information over time.

Finally, this work introduces a novel method, intersubject RSA, for measuring individual differences in neural responses using complex, naturalistic stimuli. This method and type of stimuli can provide important insights into the processing of complex social information that leads to the creation of a shared reality and facilitates social communication (49, 50). Future applications of this approach could enable us to delineate the development of high-level social cognitive abilities in the DMN during childhood, as well as to understand the development of cross-modal representations in the DMN. This method may also enable the detection of abnormalities during complex naturalistic perception and narrative interpretation relevant to psychotic disorders. For example, previous work has shown that individuals with autism spectrum disorder or schizophrenia show atypical interpretations of more simple shape-based animations (29, 51–53), but differences in interpretation under naturalistic conditions have not yet been linked to individual differences in neural responses.

In conclusion, this work provides evidence that shared understanding results in shared neural responses within and across forms of communication. The similarity between neural patterns elicited by similar interpretation of the same narrative communicated in different forms (shapes versus words) demonstrates the remarkable modality invariance and strong social nature of the default mode network. This work invokes the role of the default mode network in representing subtle differences in interpretation of complex narratives.

## Methods

### Subjects

Seventy-six adult subjects (ages 18-35, mean 22.2 years; 27 male) with normal hearing and normal or corrected-to-normal vision participated in the experiment. Three subjects were excluded for excessive motion during scanning (>3mm), one for falling asleep, and one due to an anomalous finding, resulting in 36 subjects for the Movie group, 18 for the ToM Audio group, and 17 for the Phys Audio group. A larger sample size was selected for the Movie group in order to better detect individual differences in narrative interpretation. Sample sizes were otherwise chosen based on power analyses on intersubject analyses (54). All experimental procedures were approved by the Princeton University Internal Review Board, and all subjects provided informed, written consent.

### Stimuli and experimental design

Subjects were split into three separate groups. The “Movie” group was scanned in fMRI while watching a 7-min animated film. The movie depicted a short story using moving geometric shapes in the style of Heider & Simmel (1). The narrative included complex and abstract events, such as a child going to sleep and having a dream, birds changing into a monster, and making new friends. While there was no spoken dialogue, the animation included an original piano score that communicated mood and was congruent with events in the narrative.

Two additional groups of subjects were also scanned in fMRI while listening to two different audio descriptions of the animation. The “Theory of Mind (ToM) Audio” group listened to a 7-min spoken version of the animation that interpreted the shapes as animate characters (Fig 1A, middle). The interpretation was based on the director's interpretation of the animation. The “Physical (Phys) Audio” group listened to a 7-min audio track that described the physical characteristics and motion of the shapes in the Movie without attributing animacy to them (Fig. 1A, right). Both audio descriptions were edited to be the same length as the Movie and so that the onset of events in the audios were time-locked to the onset of events in animation, following (9) and (13) (Fig. 1A). Subjects listening to the audio stimuli did not view any visual stimuli.

To remove transient, non-selective responses that occur at the onset of a stimulus, all scans were preceded by the same, unrelated 37-second movie clip. This clip was cropped from all analyses. Following stimulus presentation, subjects were asked to free recall the stimulus using their own words and in as much detail as possible (Fig. 1B).

### Stimulus presentation

Details are provided in SI Methods, Stimulus presentation.

### MRI acquisition

Details are provided in SI Methods, MRI acquisition.

#### Behavioral data analysis

Subject recalls were assessed for similarity to each other using Latent Semantic Analysis (LSA), a statistical method for representing the similarity of texts in semantic space (21). Details are provided in SI Methods, Behavioral data analysis.

#### MRI data analysis

##### Preprocessing

MRI data were preprocessed using FSL 5.0 (FMRIB, Oxford) and custom Matlab scripts. Results were visualized on 3D inflated cortical masks using NeuroElf. Details are provided in SI Methods, Preprocessing.

##### Audio correlations between stimuli

The two audio recordings and the Movie were aligned in time such that the start of each event occurred at the same time across stimuli. As a result, the audio envelops (audio amplitude) of the three stimuli may be correlated. To control for this low-level similarity across stimuli in cross-modal analyses, we followed (13) and regressed out the audio envelope from each subject’s BOLD response. Details are provided in SI Methods, Projection of audio envelops.

##### Representational similarity analysis (RSA)

To identify regions of the brain where greater recall similarity predicts greater neural similarity within and across conditions, we conducted a representational similarity analysis (RSA) (20) between recall similarity, as measured by LSA, and neural similarity, as measured by correlating response timecourses between subjects. RSA was conducted both voxel-wise over the entire brain and within independently-defined ROIs. Statistical significance was assessed using a permutation test. We corrected for multiple comparisons by controlling the False Discovery Rate (FDR) (48) of the RSA map using q criterion = 0.05. Details are provided in SI Methods, Representational similarity analysis.

##### Intersubject correlation (ISC) analysis across modalities

In addition to RSA, we conducted an intersubject correlation analysis (ISC) (22) to search for multimodal responses which are shared across subjects who watched the Movie and subjects who listen to the Audios. ISC was calculated by taking the average correlation between each Movie subject’s response timecourse to the average of either the ToM Audio subjects or the Phys Audio subjects. Statistical significance of ISC was assessed using a bootstrapping procedure based on phase randomization. To control for multiple comparisons, the family-wise error rate was controlled at q = 0.05. Details are provided in SI Methods, Intersubject correlation.

##### Default mode network ROI analysis

All analyses were also conducted on independently defined regions-of-interest (ROIs) of the DMN. Details are provided in SI Methods, DMN ROI definition.

## Author contributions

MN, TV, and UH developed the study concept, stimulus creation, and study design. Data collection and analysis was performed by MN. All authors contributed writing to the manuscript, and all authors approved the final version of the manuscript for submission.

## Acknowledgements

We would like to thank Adele Goldberg, Janice Chen, and Chris Baldassano for helpful advice on the analysis and Amy Price for helpful comments on the paper. This work was supported by NIH Grant 5DP1HD091948-02 (UH), NIH Grant RO1 (MN), and The Allison Family Foundation (TV). The animation was created by visual artist Tobias Hoffman and TV, the original score was written by Jodi S. van der Woude, and the audios were recorded by Max Rosmarin. The movie, titled “When Heider Met Simmel,” is available for research use by contacting the corresponding author.

## Supporting information

### Methods

#### Stimulus presentation

Stimuli were presented using MATLAB (MathWorks) and Psychtoolbox (56). Video was presented by LCD projector on a rear-projection screen mounted in the back of the scanner bore and was viewed through a mirror mounted to the head coil. Audio was played through MRI-compatible insert earphones (Sensorimetrics, Model S14). Subject recalls were recorded using a customized MR-compatible recording system with online sound cancelling (Optoacoustics Ltd, FOMRI II).

#### MRI acquisition

Subjects were scanned in a 3T Magnetom scanner (Prisma, Siemens) located at the Princeton Neuroscience Institute Scully Center for Neuroimaging using a 64-channel head-neck coil (Siemens). In the Audio and Movie scans, volumes were acquired using a T2*-weighted multiband EPI pulse sequence (TR 1500 ms; TE 39 ms; voxel size 2x2x2mm; flip angle 55°; FOV 192x192 mm^2^, multiband acceleration factor 4) with whole-brain coverage. Following functional scans, a fieldmap (mean and phase) was collected (dwell time 0.93 ms; TE diff 2.46 ms). Finally, a high-resolution anatomical image was collected using a T1-weighted MPRAGE pulse sequence (voxel size 1x1x1 mm).

#### Behavioral data analysis

Free recalls were lightly edited to remove non-stimulus related utterances (e.g. “I don’t remember,” “I’m done,” etc.). The edited recalls were then assessed for similarity to each other within and across stimulus groups using Latent Semantic Analysis (LSA), a statistical method for representing the similarity of texts in semantic space (21). Recall similarity was measured as the cosine distance between recalls in the semantic space. Here, the semantic space was derived from the Touchstone Applied Science Associates (TASA) college reading-level corpus with 300 factors, as implemented on lsa.colorado.edu. To order subjects by similarity to each other, we then conducted agglomerative hierarchical clustering with complete-linkage on the LSA similarity matrices.

#### Preprocessing

MRI data were preprocessed using FSL 5.0 (FMRIB, Oxford), including 3D motion correction, fieldmap correction, linear trend removal, high-pass filtering (140 Hz), and spatial smoothing with a Gaussian kernel (FWHM 4 mm). All data was aligned to standard 2-mm MNI space. Following preprocessing, the first 60 TRs were cropped to remove the introductory videos and transitory changes at the start of the stimulus. Voxels with low mean signal (2 std below average) were also removed. Data was z-scored over time. All analyses were conducted in volume space using custom Matlab scripts and then visualized on 3D inflated cortical masks using NeuroElf (http://neuroelf.net).

#### Projection of audio envelops

Because the Movie and two Audios were aligned in time such that the start of each event occurs at the same time across stimuli, the audio envelops (audio amplitudes) may be correlated. Following (13), for between-condition analyses, we thus projected out the audio envelop from each subject’s neural response. The audio envelop for each stimulus was calculated using a Hilbert transform and then down-sampled to the 1.5-second TR using an antialiasing, low-pass finite impulse response filter. The resulting envelops were then convolved with a hemodynamic response function (57). The envelops were entered into a linear regression model for each voxel in each subject in the corresponding condition. For between-condition analyses, the BOLD response timecourse was then replaced with residuals of the regression.

#### Representational similarity analysis

To identify regions of the brain where greater recall similarity predicts greater neural similarity within and across conditions, we conducted a representational similarity analysis (RSA) (20) between LSA recall similarity and intersubject neural correlations. First, a matrix of neural similarity between every pair of subjects within or between conditions was calculated for each voxel by correlating each subject’s response timecourse with every other subject’s response time course. Spearman’s r was then calculated between this matrix of neural similarity and the matrix of LSA recall similarity (Fig. 2). This analysis was conducted both within the Movie group and between the Movie group and each Audio group. For the within group RSAs, the neural and recalls similarity matrices are symmetrical, so only the lower triangles are correlated. For the between group RSAs, the entire matrix is correlated. This analysis was restricted to gray matter voxels of the brain.

Following (20), statistical significance for RSA was assessed using a permutation test. For each voxel, the rows and columns of the neural similarity matrix were randomly shuffled, and the resulting shuffled matrix was correlated with the LSA similarity matrix as described above. This shuffling procedure was repeated 1000 times, resulting in a null distribution of 1000 values for the null hypothesis that there is no relationship between recall similarity and neural similarity. The mean and standard deviation of the null distributions were used to fit a normal distribution and calculate p-values. We corrected for multiple comparisons by controlling the False Discovery Rate (FDR) (55) of the RSA map using q criterion = 0.05.

#### Intersubject correlation

In addition to RSA, we conducted an intersubject correlation analysis (ISC) (2, 22) to search for cross-modal responses which are shared across subjects who watched the Movie and subjects who listened to the Audios. ISC was calculated by correlating each movie subject’s response timecourse to the average timecourse of ToM Audio or Phys Audio subjects in the same voxel. The average of these correlations across subjects is taken as the ISC. To compare shared responses between the movie group and the ToM audio group, as a function of interpretation, we calculated the average LSA similarity between each Movie subject and the ToM Audio subjects. We then took the 18 Movie subjects who were most similar behaviorally to the ToM Audio group and calculated ISC between them, and then repeated the analysis on the 18 Movie subjects who were least behaviorally similar to the ToM Audio group. We conducted the same procedure for comparing the Movie subjects and the Phys Audio subjects.

Statistical significance of ISC was assessed using a permutation test. Following (9), each voxel’s time course was phase-scrambled by taking the Fast Fourier Transform of the signal, randomizing the phase of each Fourier component, and then inverting the Fourier transformation. This randomization procedure thus only scrambles the phase of the signal, leaving its power spectrum intact. Using the phase-scrambled surrogate dataset, the ISC was again calculated for all voxels as described above, creating a null distribution of average correlation values for each voxel. This bootstrapping procedure was repeated 1000 times, producing 1000 bootstrapped correlation maps.

To correct for multiple comparisons, the largest ISC value across the brain for each bootstrap was selected, resulting in a null distribution of the maximum noise correlation and representing the chance level of calculating high correlation values across voxels in each bootstrap. The family-wise error rate of the measured maps was controlled at q = .05 by selecting a correlation threshold (R*) such that only 5% of the null distribution of maximum correlation values exceeded R*. In other words, only voxels with mean correlation value (*R*) above the threshold derived from the boot-strapping procedure (*R*\*) were considered significant after correction for multiple-comparisons and were presented on the final map.

#### DMN ROi definition

In addition to whole-brain, voxel-wise analyses, RSA and ISC analyses were also conducted on independently-defined ROIs. These ROIs were defined using functional connectivity on previously published, independent data (10) where subjects were scanned in fMRI watching a movie. A seed ROI for posterior medial cortex was taken from a resting state-state connectivity atlas (posterior medial cluster functional ROI in “dorsal DMN” set) (59). Following (10), the DMN ROIs were then defined by correlating the average response in the PMC ROI to every other voxel in the brain during the movie for each of 17 subjects, averaging the resulting connectivity map, and thresholding at R = 0.5 (Fig. 1C). Although the DMN is typically defined using resting-state data, recent work has shown that the same network is activated during temporally extended stimuli (8).

### M1: Animated shapes movie

Animated shapes movie, “When Heider Met Simmel.” Written, produced, and directed by Tamara Vanderwal. Animation by Tobias Hoffman. Original score written by Jodi S. van der Woude.

